# Controlling treatment toxicity in ovarian cancer to prime the patient for tumor extinction therapy

**DOI:** 10.1101/2025.07.10.664235

**Authors:** Kit Gallagher, Rachel S. Sousa, Chandler Gatenbee, Ryan Schenck, Peng Chen, Timon Citak, Sydney Leither, Lucia Mazzacurati, Agata Xella, Zoe Zhou, Dawn Lemanne, Paulo Rodriguez, Erin George, Maximilian A. R. Strobl

**Affiliations:** Wolfson Centre for Mathematical Biology, Mathematical Institute, Oxford, UK; Department of Integrated Mathematical Oncology, H. Lee Moffitt Cancer Center, Tampa, Florida, USA; Center for Complex Biological Systems, University of California Irvine, Irvine, California, USA; Stanford Cancer Institute, Stanford University, California, USA; Department of Translational Hematology and Oncology Research, Cleveland Clinic, Ohio, USA; Department of Physics, Case Western Reserve University, Ohio, USA; Max-Planck-Zentrum für Physik und Medizin & Max Planck Institute for the Science of Light, Erlangen, Germany; Department of Physics, Friedrich-Alexander-University Erlangen-Nürnberg, Erlangen, Germany; Department of Computer Science and Engineering, Michigan State University, Michigan, USA; Department of Gynecologic Oncology, H. Lee Moffitt Cancer Center, Tampa, Florida, USA; Oregon Integrative Oncology, Ashland, Oregon, USA; Department of Medicine, University of Arizona, Tucson, Arizona, USA; Department of Immunology, H. Lee Moffitt Cancer Center, Tampa, Florida, USA

**Keywords:** Ovarian Cancer, Mathematical Modeling, Extinction Therapy, Toxicity Management

## Abstract

High-grade serous ovarian cancer (HGSOC) remains a major clinical challenge. In particular among those patients with homologous recombination (HR)-proficient tumors (*>*50%), most eventually succumb to their disease due to high recurrence rates, acquired resistance, and cumulative toxicity. This report summarizes work from the 12^th^ IMO Workshop in which we explored an alternative “extinction therapy” strategy for frontline treatment of HGSOC. Inspired by ecological principles, this multi-strike approach aims to eradicate tumors not through a singular “magic bullet” but through a series of therapies after standard frontline treatment when the tumor is still, and perhaps most, vulnerable. We present a framework leveraging mathematical modeling (MM) to develop personalized multi-strike protocols for HGSOC. Key contributions include: 1) An “IMOme” score using liquid biopsy data to assess patient-specific hematopoietic toxicity risk, guiding the timing and selection of subsequent therapies, 2) MM strategies to design effective lowdose combinations of targeted agents to achieve synthetic lethality while managing toxicity, and 3) A MM framework to analyze the interplay between chemotherapy, gut microbiome toxicity, and immunotherapy, demonstrating how mitigating microbiome damage could enhance immune response. Overall, the computational approaches presented herein aim to support the design of personalized, multi-strike regimens in the frontline setting that proactively target tumor extinction while managing toxicity, ultimately seeking to deliver cures for patients with HGSOC.

## 1. Introduction

High-grade serous ovarian cancer (HGSOC) remains the most lethal gynecologic malignancy. While frontline surgery and chemotherapy lead to complete remission in 87% of patients, 82% of these recur and ultimately succumb to their disease [1]. Not only do recurrences become progressively more difficult to treat due to drug resistance, but cumulative toxicity from repeated chemotherapies and worsening disease burden limits the ability to provide subsequent therapies. Thus, there is an urgent need to innovate not only *what* drugs we give but also *when* and *how* we use them to increase chances for cure.

This report summarizes key findings from our group project at the 12^th^ IMO Workshop: Toxicity, held in Tampa, FL, between 4^th^ and 8^th^ November 2024, where we investigated an *extinction therapy* approach to HGSOC treatment. The current HGSOC treatment paradigm is based on the maximum tolerable dose (MTD) principle, which seeks to maximize the chance of cure by administering therapy at the highest tolerable dose and frequency to maximize tumor cell kill. However, when considering the entirety of the patient’s cancer there is a multiplicity of resistance mechanisms and protected ecological niches that cells may occupy making for a diverse tumor cell population that can acclimate and adapt to toxic pressures. Thus, despite initial responses our traditional treatment approaches fail because of a mismatch between our evolutionary static treatment approach and cancer’s ecological and evolutionary dynamics [2]. In contrast, multi-strike extinction therapy proposes to use a series of “strikes” that drive the tumor to extinction by both killing tumor cells and modifying the tumor environment to gradually eradicate any surviving cancer cells [3, 4].

The concept of “extinction therapy” was introduced by Gatenby et al. [3, 4] and is based on the observation that natural extinctions rarely happen as the result of a singular catastrophic event. Instead, extinctions typically occur due to a sequence of events that combine to inflict large and persistent population losses as well as durable environmental change. Populations are most vulnerable when small, due to the risk of random extinction and their reduced diversity (genetic, phenotypic, and geographic) - the key ingredient for Darwinian evolution. Multi-strike extinction therapy thus postulates that to maximize the chance of cure, we should focus not on maximizing the kill achieved by any particular line of therapy, but by pursuing a sequence of treatments that target different population vulnerabilities and that can be sustained even once - and especially once - the tumor is small.

We present a framework to translate these eco-evolutionary principles to the treatment of HGSOC. Specifically, we explore how to combine frontline platinum-based chemotherapy with a “multi-strike consolidation phase” that consists of a series of carefully tailored “second strike” therapies (Figure 1). These therapies extend the duration of disease control and maximize the chance of cure before cumulative toxicity and drug resistance curtail therapeutic options. Recently, the potential of a well-designed multi-strike approach in HGSOC has been demonstrated in the subgroup of patients with homologous recombination (HR)-deficient disease. HR deficiency can be exploited therapeutically by PARP-inhibitors (PARPis), but the success of PARPis as single agents has been limited. However, their introduction as maintenance therapy following frontline chemotherapy - i.e. as a second strike that follows chemotherapy - has revolutionized the care of HR-deficient patients, particularly those with *BRCA1/2* mutations, achieving improvements in progression free survival of 28 months in HR-deficient patients with BRCA mutations (HR 0.38, 95% CI 0.32-0.46, p*<* 0.001) and 9.2 months in HR-deficient patients without BRCA mutations (HR 0.47, 95% CI 0.34-0.65, p*<* 0.001)[5]. In BRCA1/2 patients at the 7-year follow-up, 67% of olaparib patients versus 46.5% of the placebo patients were alive and 45.3% versus 20.6% were alive and had not received first subsequent treatment [6]. This success story raises the question: how do we extend the multi-strike paradigm beyond patients with *BRCA1/2* mutations, especially the HR-proficient subgroups (*>* 50% of patients), who have seen no improvement in long-term survival for decades, and represent an important, unmet clinical need?

**Figure 1:**
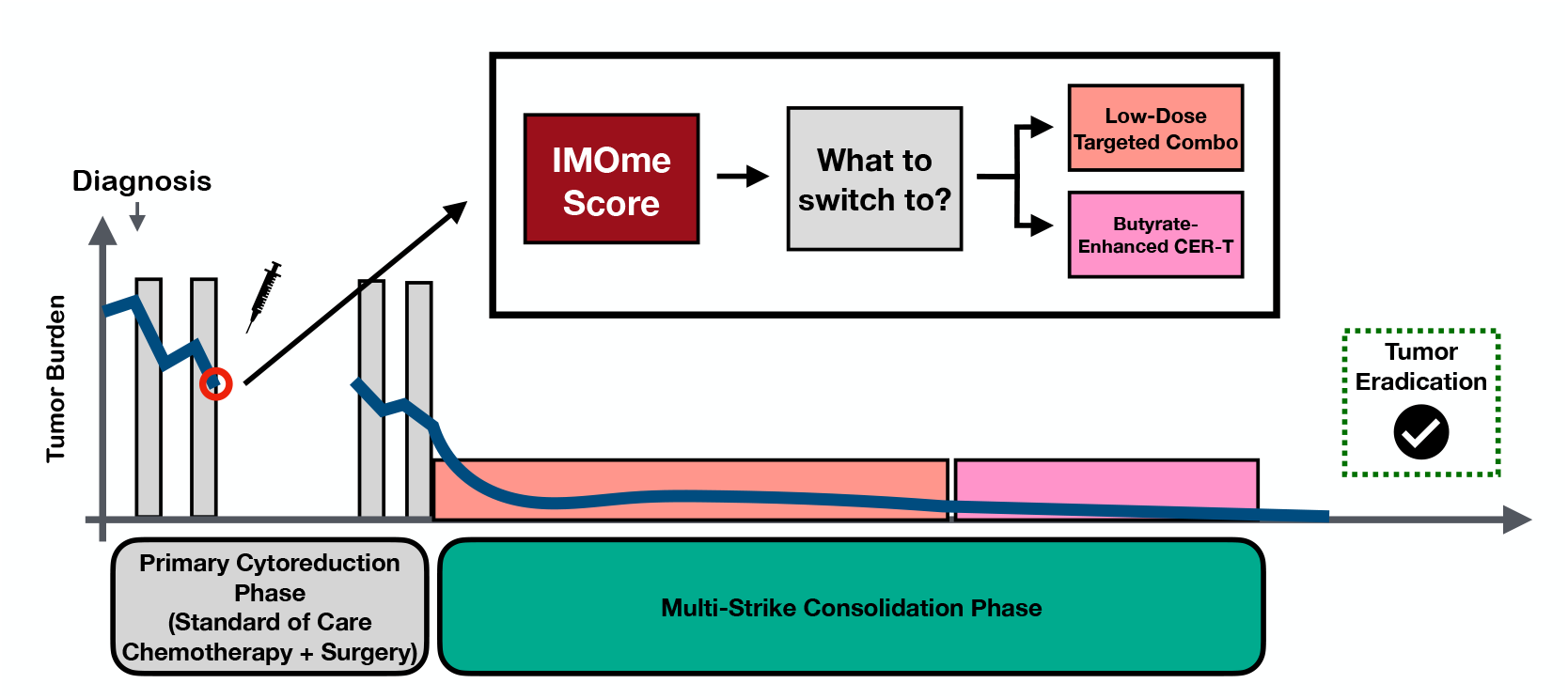
We propose that to maximize the chance of curing patients with HGSOC we require multi-strike protocols that eradicate the tumor through a series of treatments (“strikes”) during first-line treatment. The key challenges in developing these protocols are dose-limiting toxicity and the limited numbers of therapeutic options. In our IMO workshop project, we developed three approaches to tackle these issues: i) a novel methylation-based IMOme score to stratify patients according to risk of hematopoetic toxicities, and mathematical modeling to support the development of second strike options during the consolidation phase in the form of (ii) low-dose combination regimens and (iii) approaches to reduce gut microbiome toxicity and potentiate immunotherapy.

In the following, we discuss how to leverage liquid biopsy data, machine learning and mathematical modeling to support the development of multi-strike protocols for HGSOC (Figure 1). Specifically, we address two critical questions in the design of these protocols: i) when to switch treatment agents, and ii) which agent(s) to switch to? In Section 2.1, we present a machine learning classifier to identify patients at high-risk of hematologic toxicity from first-line therapy, informing when to switch therapy and what type of second strikes may be tolerated. Subsequently, we discuss two approaches for the development of second strikes to overcome the lack of singular targetable mutations in this group of diseases: i) low-dose combinations of small molecule inhibitors targeting cell cycle regulators, and ii) immunotherapy. We hypothesize that despite limited molecular targets and the challenges of dose-limiting toxicities, strategic combinations and sequencing of small molecule inhibitors can simultaneously perturb multiple pathways and create a state of synthetic lethality from individually tolerable inhibitions, thereby effectively killing cancer cells. In Section 2.2, we build on emerging work on the combination of PARP with WEE1 or ATR inhibitors and demonstrate how mathematical modeling can help to overcome the combinatorial complexity of choosing when and how to best combine these agents. Finally, in Section 2.3, we discuss how to increase the efficacy of immunotherapy in HGSOC by leveraging the gut microbiome and its metabolic products, which have emerged as central mediators of anti-tumor immunity. Toxicity to the gut microbiome is ubiquitous during cancer therapy yet has remained largely unaddressed. We present a novel mathematical model which enables us to systematically explore these dynamics and we show proof-of-principle simulations of how mitigating microbiome toxicity through therapeutic supplements or earlier cessation of chemotherapy could significantly potentiate adjuvant immunotherapy. Our long-term goal is to create innovative strategies that effectively predict, prevent, and overcome toxicity and leverage a series of targeted therapeutic approaches that maximize chances for cure for patients with HGSOC, particularly HR-proficient patients, in the frontline setting.

## 2. Results & Discussion

### 2.1 Assessment of bone marrow health: the IMOme score

A key challenge in the treatment of HGSOC is the gradual build-up of treatment-inflicted damage to the patient’s bone marrow (hematologic toxicity). Resulting complications, such as anemia, neutropenia, and thrombocytopenia, eventually prevent the administration of further therapy, even when effective options are available. This limitation on the total amount of tolerable therapy underscores the importance of carefully selecting the timing and type of treatment. Therefore, to design multi-strike therapies for HGSOC it is crucial to develop methodologies that can predict a patient’s tolerance to myelosuppressive therapy, so that the most effective combination and sequence of drugs can be chosen for a particular patient (or subgroups of patients). For example, if a patient is classified as high risk, we might consider second strike options that are less myelosuppressive (e.g., bevacizumab) or even reduce the number of upfront chemotherapy cycles to allow for more second strike options in the consolidation phase.

To address this challenge, we investigated emerging liquid biopsy technologies to assess bone marrow resilience. We developed the *IMOme* metric which characterizes the diversity of peripheral blood cell subsets using “fluctuating methylation clocks” [10]. DNA methylation, the attachment of methyl groups to DNA particularly at CpG sites (regions where cytosine is followed by guanine), is a key mechanism for controlling gene expression. However, methylation also occurs reversibly and stochastically, at a remarkably constant average rate in regions which are not actively involved in gene expression [11]. This provides a molecular clock to track cell ancestry: the more similar the methylation patterns between two cells, the shorter the time since their last common ancestor (Figure 2a). Note that this is in contrast to the approach of conventional methylation-based molecular clocks, which count the irreversible accumulation of methylation changes since birth to infer the relatedness of lineages over much longer timescales [12].

**Figure 2:**
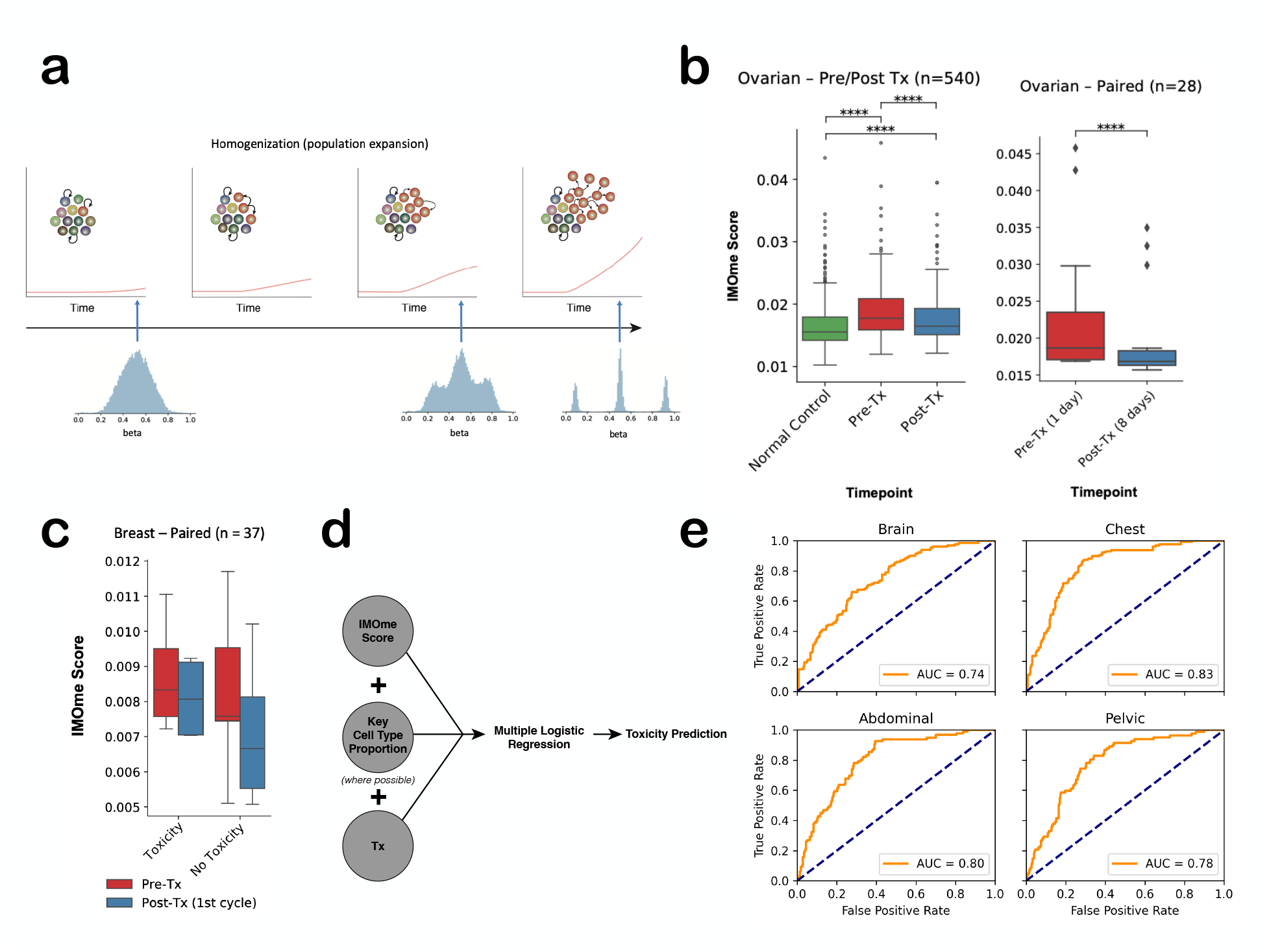
**(a)** The IMOme score to estimate bone marrow resilience from bulk methylation data. As stem cells divide, the resulting lineages accumulate diverging methylation patterns in non-coding regions of the genome due to random methylation changes. This provides a molecular clock to characterize stem cell diversity. **(b)** The IMOme score is significantly reduced after chemotherapy treatment, in both un-paired [7] and paired [8] datasets. **(c)** The reduction is IMOme score is lessened in patients with reported toxicities (based on cardio-toxicity in breast cancer treatment [9]), enabling early prediction of future toxicities before they are clinically measurable. **(d)** Using a multiple logistic regression model, we can predict with high accuracy which patients experience specific toxicities induced by prior cancer treatment exposure across a range of cancers, based on post-treatment methylation samples. **(e)** ROC curves for toxicity prediction from each region-specific radiotherapy dose, with the labeled AUC (area under curve) showing a high degree of predictive accuracy.

Gabbutt et al [10] recently developed a computational method to infer the diversity and activity of the hematopoietic stem cell pool from bulk methylation data extracted via standard methylation arrays from a simple blood draw. Here, we leveraged this method to develop the “IMOme Score” which quantifies the ability of the bone marrow to regenerate after chemotherapy. Specifically, the score characterizes clonal diversity of the hematopoietic stem cell population, with a low score indicating a clonally homogeneous population derived from a very recent common ancestor (Figure 2a). We hypothesize that low diversity may correspond to severely depleted hematopoietic stem cell populations and identify patients at higher risk of bone marrow toxicity.

To begin with we tested whether we can detect changes in the IMOme score following chemotherapy. We contrast methylation data from patients with epithelial ovarian cancer pre/post treatment (N = 266) with an age-matched healthy population (N = 274) [8]. We observe a significantly higher IMOme score across cancer patients (pre and post treatment) relative to the comparison cohort, indicating elevated clonal diversity in patients with cancer (Figure 2b). Moreover, in accordance with our hypothesis, we find that the IMOme score was significantly reduced post therapy compared to pre therapy levels. Furthermore, considering paired pre/post treatment samples from a smaller cohort (N = 14) [7], we were able to detect a significant reduction in the IMOme score only 3 days after the end of the first chemotherapy cycle (5 days of Decitabine; Figure 2b). Together, these observations suggest that the diversity of the hematopoietic stem cell pool decreases during chemotherapy.

Next, we validated IMOme’s predictive power for future toxicities in a breast cancer cohort (N = 19) with paired samples (pre/post treatment) and recorded cardio-toxicities ([9]; Figure 2c). Patients developing cardio-toxicities showed a smaller post-treatment IMOme decrease, suggesting impaired hematopoietic stem cell restoration. Notably, IMOme predicted this toxicity after just one cycle of chemotherapy, while routine monitoring via echocardiograms only detected it after four cycles. These results highlight IMOme’s potential for early identification of at-risk patients, enabling timely treatment modifications to avoid inflicting additional toxicity that could have been prevented after the first cycle.

Finally, we investigated how the IMOme score can be further improved by integrating additional data modalities. Bulk methylation data from also allows deconvolution of circulating immune cell type distributions. Applying this method to bulk methylation data collected post-treatment from a large pediatric cancer cohort (N = 968) [14], we found increased natural killer cell, monocyte, CD4 memory T-cell, and neutrophil populations among patients with reported toxicity. Integrating these changes into a multivariate logistic regression model with IMOme accurately predicted a range of adverse toxicity events associated with prior cancer treatment exposure (Figure 2d). We validate our toxicity prediction against long-term cardiometabolic toxicities from region-specific radiotherapy doses across this cohort, with an average AUC of 80% (Figure 2e).

In summary, we have shown that we can identify systematic changes in the clonal and cell type distributions of the peripheral blood. Using this metric, we develop a predictor of future toxicity. This informs future treatment decisions, including second line drug choice and scheduling for extinction therapy.

### 2.2 Mathematical cell cycle modeling as a guide to low-dose combination therapy optimization

The effectiveness of PARPi maintenance in HR-deficient, particularly *BRCA1/2* mutant, patients underscores the promise of multi-strike strategies to HGSOC. A key obstacle in extending this concept to HR-proficient patients is the absence of readily apparent therapeutic targets for a “second strike”. However, recent work by us and others has shown that multiple cell cycle checkpoint inhibitors can synergize to induce synthetic lethality where individual inhibitors do not provide sufficient effect [15, 16]. We demonstrated that PARPi in combination with ATR inhibition (ATRi) resulted in synergy and complete and durable therapeutic responses that significantly increased survival in platinum and acquired PARPi-resistant patient-derived xenograft models [15]. We also showed that WEE1 and ATR inhibition was a synergistic drug combination for Cyclin E1 overexpressing HGSOC [16]. Yet, the sheer number of potential combinations and schedules poses a significant practical hurdle. For example, with just two drugs with two dose levels each and four weeks to a treatment cycle there are 4^4^ = 256 possible schedules to evaluate.

Recently, mathematical modeling has established itself as a powerful way to tackle this complexity, and to rapidly and cheaply shortlist candidates for empirical testing [17, 2]. However, this approach requires carefully calibrated and validated mathematical models, of which there exist few for HGSOC [18]. To address this gap, we worked on mathematical modeling to predict and optimize a sequence of targeted therapy combinations for HGSOC to minimize toxicity and improve efficacy for PARP and ATR inhibitors. We adopted a model of cell cycle progression developed by Pugh et al [13], which categorizes tumor cells in G1, S, S damaged (SD), G2/M, and G2/M damaged (G2D) states and uses ordinary differential equations (ODEs) to describe how cells transition between different compartments over time (Figure 3a; see 4.2 for details). In the absence of treatment, cells follow the regular progression of G1 *→* S *→* G2/M *→* M *→* S. To represent the effect of PARP inhibition, which prevents the repair of single stranded DNA breaks, the model assumes that affected cells are pushed from G1 phase into a damaged SD state rather than to S phase. PARPis also inhibit DNA repair of a G2D state, preventing cells from moving to G2/M phase. ATRi functions in two ways in the model: i) ATRi inhibits the repair of DNA damage in SD and ii) ATRi can also result in the abrogation of the G2/M checkpoint, which subsequently causes cells to undergo mitotic catastrophe (Figure 3a).

**Figure 3:**
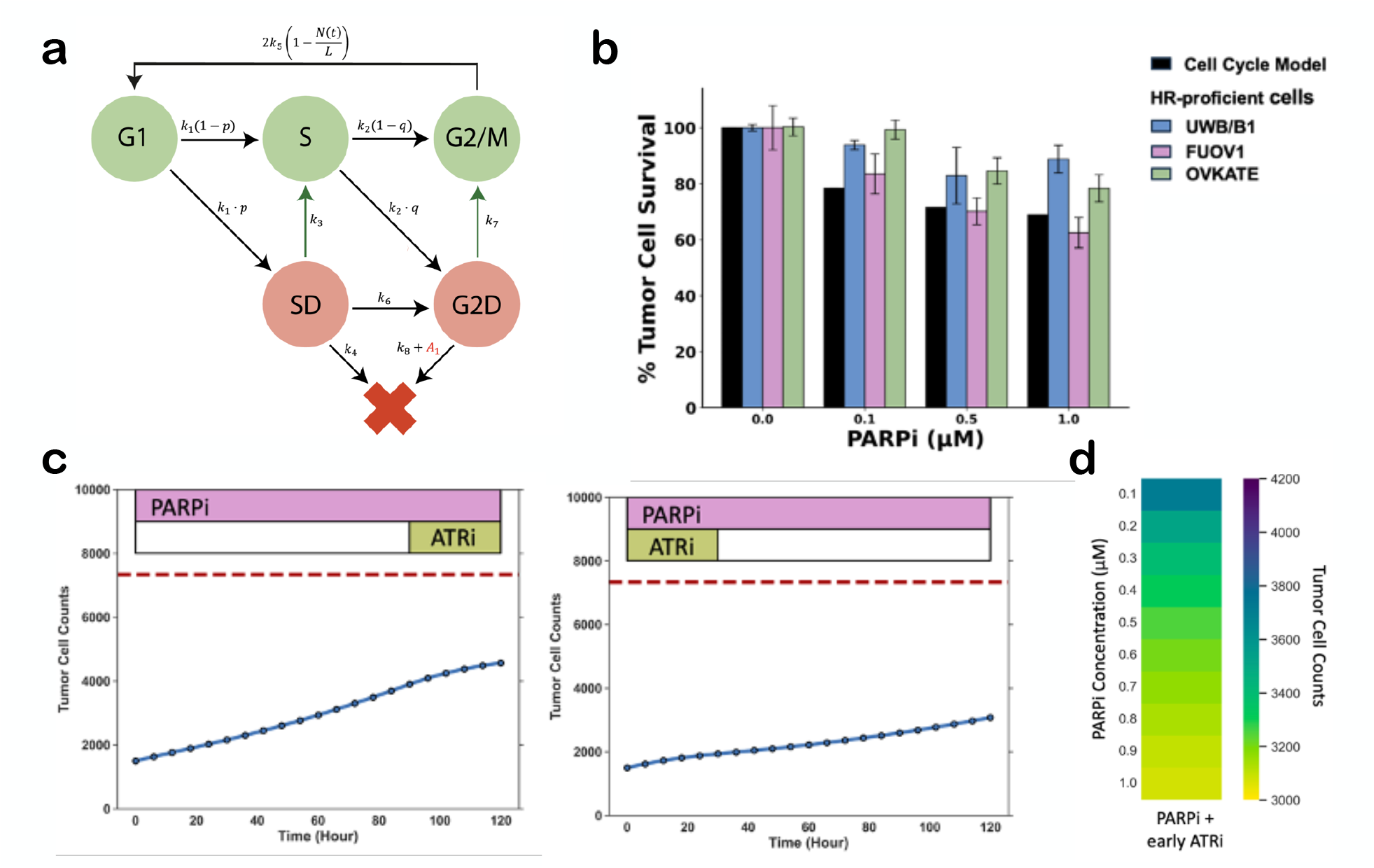
**(a)** The cell cycle mathematical model by Pugh et al [13] (figure reproduced from ref [13]). **(b)** Cell cycle model compared to ovarian cancer cell lines treated with PARPi, showing that the model can recapitulate the behavior of HGSOC cells *in vitro*. **(c)** Combination PARPi + ATRi where ATRi is given at different time points, showing that early administration of ATRi is favored (each drug is dosed at 1*µ*M). Red dashed line: number of tumor cells without treatment at hour 120. Blue line: tumor cell counts with specified treatment. **(d)** Lower concentrations of PARPi when given with early ATRi show similar final tumor cell counts (ATRi dosed at 1*µ*M).

Next, we validated that the model could recapitulate published *in vitro* drug response data in ovarian cancer cell lines [15]. Given the time constraints of the workshop, we adopted parameter values by Pugh et al [13] as a proof-of-principle. Even though these values were derived from a head and neck cancer cell line (FaDu ATM-KO cells), we found that the so-parameterized cell cycle model predicted a similar percentage of tumor cell survival to three HR-proficient ovarian cancer cell lines (UWB/B1, FUOV1, and OVKATE) when treated with various doses of PARPi and ATRi monotherapy (Figure 3b). The model predictions also qualitatively re-capitulated the levels of synergy of the combination of PARP and ATRis observed *in vitro* in the ovarian cancer cell lines (not shown).

Next, we explored how the model could support optimal dosing and scheduling strategies. As a first step, we compared high-dose monotherapy of PARPi or ATRi with a low-dose combination regimen of both agents. Our simulations predicted that the combination is more effective at depleting tumor cells than the monotherapies despite the lower dose levels (not shown). This prediction agrees with experimental observations by Kim et al [15], thereby further validating the model. These results could be clinically relevant, as lower doses of these inhibitors could lead to less toxicity for patients.

Subsequently, we considered how the sequence and dose in which the drugs are administered in the combination regimen shapes therapy outcome. We found in our simulations that when giving a combination of PARPi and ATRi treatment, early administration of the ATRi leads to the highest tumor cell death (Figure 3c). Furthermore, by using the model to exhaustively explore the range of possible combinations of dose levels and treatment timings, we predicted that when giving a combination of PARPi with early ATRi treatment, we can further lower the dose of the PARPi without significant loss of efficacy. Specifically, a PARPi dose of 1 *µ*M is predicted to only eliminate 2.4% more tumor cells than a PARPi dose of 0.5 *µ*M (Figure 3d). This result is interesting as early ATRi administration plus PARPi is currently being tested in clinical trials [19, 20]. Our findings suggest that we could further lower the dose of the PARPi to lower the toxicity to patients while still achieving similar tumor clearance.

### 2.3 Priming the patient for immunotherapy by mitigating chemotherapy-induced toxicity to the gut microbiome

The human gut microbiota and their metabolites play a pivotal role in maintaining physiological homeostasis, and the interplay with anti-cancer therapies is increasingly recognized as a determinant of therapeutic success [21, 22]. In particular, the production of short-chain fatty acids (SCFA) by a healthy gut microbiome is positively associate with treatment response [23]. The reason behind the importance of SCFAs is thought to be their key immunomodulatory role. SCFAs enhance immune responses by promoting the activity of cytotoxic T lymphocytes (CTLs), thereby boosting antitumor reactivity [24]. However, cancer treatments, such as carboplatin/paclitaxel chemotherapy, PARPis, and targeted therapies for ovarian cancer are associated with cumulative microbiota damage [25], and there is evidence that it does not recover without intervention [25]. We hypothesize that chemotherapy-inflicted microbiota damage is a key factor limiting the success of immunotherapy in the treatment of HGSOC by compromising immune-mediated antitumor activity and fostering an immuno-suppressive microenvironment. We propose that addressing this disruption prior to adjuvant immunotherapy may improve treatment outcomes and add a crucial further pillar to a multi-strike extinction strategy.

To explore the impact of chemotherapy on the microbiome and in-turn on the endogenous immune system we developed a simple mathematical model. Our model assumes that cancer cells grow exponentially and are predated on by the immune system at rates that are modulated by the gut microbiome, where *C*(*t*) and *I*(*t*) represent the densities of tumor and immune cells as functions of time, *t*, respectively. For simplicity, we do not explicitly model the microbiome but instead focus on *P* (*t*), the ratio of long to short chain fatty acids produced by the microbiome. This ratio has been shown to be a key modulator of immune state, shifting the immune environment from anti- to pro-tumor activity as *P* increases from 0 to 1 (corresponding to a relative depletion of SCFAs; [26, 27]). Correspondingly, we assume in our model that values of *P* near 0 (corresponding to high levels of SCFAs) promote tumor cell killing by immune cells, whereas values of *P* near 1 suppress tumor cell kill and promote tumor growth. Finally, we integrate the impact of chemotherapy, which we assume kills tumor cells and induces microbiome damage that shifts *P* towards 1 (see Section 4.3 for details).

Using our model, we explore the shifting tradeoff between tumor cell kill and cumulative microbiome toxicity with each cycle of chemotherapy. As shown in Figure 4a, sustained chemotherapy (x-axis) rapidly reduces tumor burden, but eventually the rate of reduction plateaus (left y-axis, in red), suggesting diminishing returns with each additional round of therapy. In contrast, microbiome toxicity (right y-axis) continues to increase with prolonged chemotherapy. These simulations reveal an important drawback of the current MTD approach to HGSOC therapy: by prioritizing short-term tumor reduction at diminishing therapeutic returns, we inflict significant cumulative microbiome toxicity - which may inadvertently curtail future treatment options.

**Figure 4:**
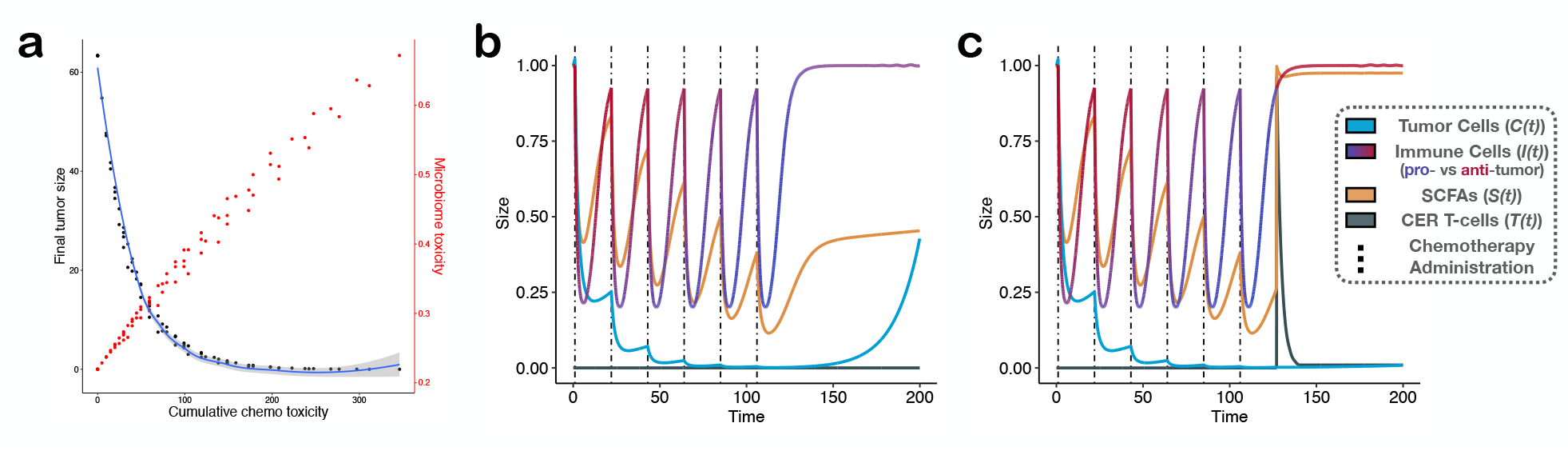
**(a)** Simulations show the diminishing returns and costs of prolonged chemotherapy, with minimal improvements in tumor reduction but increased microbiome toxicity. **(b)** Simulation of chemotherapy followed by CER T-cell injection results only in a temporary response, as destruction of the microbiome reduces SFCA, increasing immune suppression and reducing CER T-cell efficacy. Vertical lines mark chemotherapy administration. **(c)** Co-administration of SFCAs is predicted to buffer the chemotherapyinduced toxicity to the microbiome, thereby preserving a more pro-inflammatory immune environment. This increases CER T-cell efficiency and results in tumor elimination. This figure presents proof-of-principle simulations where all variables are unit-less and illustrative.

To put this observation in context, we investigated how accumulating microbiome toxicity may affect subsequent immunotherapy. Recently, the development of chimeric endocrine receptor (CER) T-cell therapy has opened up an exciting new treatment option for HGSOC. CER T-cells are engineered to target the follicle stimulating hormone receptor, which was found to be constitutively expressed on HGSOC cells, and thus these T-cells allow targeted killing of tumor cells [28] and are currently being tested in a Phase 1 clinical trial (NCT05316129). We hypothesize that chemotherapy-induced microbiome damage may reduce the efficacy of CER T-cells and that their efficacy can be enhanced by supplementing SCFAs in the patient’s diet (e.g. via addition of butyrate).

To test this hypothesis, we incorporated CER T-cells into our mathematical model and simulated how a patient would respond to the sequential administration of six cycles of chemotherapy followed by injection of one dose of CER T-cells (see Section 4.3 for details). As shown in Figure 4b, chemotherapy causes a depletion of SCFAs and an associated shift of the immune environment towards an anti-inflammatory state (*P* (*t*) *→* 1; line for *I*(*t*) changing color from red to blue). While injection of CER T-cells can induce a complete response, the immuno-suppressive microenvironment prevents tumor elimination and eventually the tumor recurs (Figure 4b). In contrast, when we co-administer butyrate with chemotherapy we can preserve a more pro-inflammatory microenvironment post-chemotherapy, thus enabling the CER T-cells to eliminate the tumor (Figure 4c).

Together, these simulations suggest that dietary SCFA supplementation, and potentially a reduction in the number of chemotherapy cycles, may be promising treatment strategies to minimize toxicity and increase CER T-cell efficacy. Such a holistic therapeutic approach is critical and novel in ovarian cancer, where standard therapies often compromise the microbiota and may limit the effectiveness of immune-based treatments. Treating the gut microbiota as an active participant in cancer therapy has significant potential to improve both the success of treatments and the quality of life for patients.

## 3. Discussion & Future Work

While significant progress has been made in treating HR-deficient HGSOC with PARPis, especially for patients with *BRCA1/2* mutations, the majority of patients have experienced few improvements in longterm survival. This substantial subgroup represents a critical unmet clinical need, demanding innovative therapeutic strategies beyond standard cytotoxic chemotherapy.

The concept of multi-strike extinction therapy, which draws inspiration from ecological principles of natural extinction events, offers a compelling paradigm shift in how we approach HGSOC treatment. The success of PARP inhibitors as maintenance in *BRCA1/2* mutant patients already hints at the power of sequential “strikes” in potentially curing HGSOC. We propose that a multifaceted and persistent therapeutic strategy, rather than solely relying on maximal initial cytotoxicity, may enable to achieve more durable responses and ultimately increase the chances of cure in this aggressive malignancy.

In this report, we have presented three steps towards a framework which integrates liquid biopsy data, machine learning, and mathematical modeling to rationally design and optimize multi-strike protocols for HGSOC (Figure 1). Our investigations into predicting bone marrow toxicity through the IMOme score suggest a novel means for personalizing the timing and selection of subsequent therapies, aiming to minimize cumulative toxicity and preserve treatment options. In parallel, our explorations of low-dose inhibitor combinations and microbiome-enhanced immunotherapy provide promising avenues for developing effective “second strike” options that can overcome the paucity of clear molecular targets leveraging already approved treatments and supplements.

The work discussed in this report was produced over the course of a 5-day workshop and primarily serves to motivate further investigation. In the next step, we plan to collect retrospective clinical data (methylation panels on PBMCs and circulating fatty acid levels) to validate our findings involving the IMOme score and calibrate our mathematical model from Section 2.3. At the same time, we also plan to pursue *in vitro* experiments to first calibrate the cell cycle mathematical model from Section 2.2 and subsequently validate our predictions. We believe that carefully calibrated and validated mathematical modeling frameworks will be invaluable in navigating the inherent combinatorial complexity of a multi-strike approach, allowing for the *in silico* prioritization of treatment schedules for future empirical validation.

Ultimately, our findings contribute to a growing body of evidence suggesting that a more nuanced and strategically sequenced approach to HGSOC treatment that moves beyond the traditional MTD paradigm holds significant promise. Extending this multi-strike philosophy to all HGSOC patients, guided by predictive biomarkers, rationally designed drug combinations, and an appreciation for the tumor microenvironment, represents a critical step forward in improving long-term outcomes for these patients. Future research will focus on translating these computational insights into clinically feasible and effective treatment protocols, with the ultimate goal of transforming HGSOC from a largely incurable to a potentially eradicable disease.

## 4. Methods

### 4.1 Biomarker Development

In Table 1 we summarize the data used in developing and validating the IMOme score. The IMOme score was calculated from the beta variance of the distribution of fluctuating CpG loci state, using a method previously published in [10]. Cell type deconvolution was done using CIBERSORT. Multivariate logistic regression was carried out using sklearn v1.5.2, with L2 regularization.

**Table 1:** Overview of cohorts utilized to test the IMOme score.

| Source | Cancer | Treatment | Toxicity | Phase | Paired | Patients |
| --- | --- | --- | --- | --- | --- | --- |
| Teschendorff et al., 2010 [8] | Ovarian | Chemotherapy | No | Pre/Post | No | 540 |
| Fang et al., 2010 [7] | Ovarian | Chemotherapy | No | Pre/Post | Yes | 18 |
| Bauer et al., 2021 [9] | Breast | Chemotherapy | Yes | Pre/Post | Yes | 19 |
| Sehl et al., 2020 [31] | Breast | Adjuvant | No | Pre/Post | Yes | 72 |
| Song et al., 2022 [14] | Pediatric | Various | Yes | Post | N/A | 2052 |

Long-term toxicity events from Song et al., 2022 [14] were extracted from medical records using a structured protocol [29], and region-specific radiotherapy (RT) dosimetry were estimated from radiation oncology treatment records [30]. We included grade 1+ events from seven common cardiometabolic chronic health conditions (CHCs): abnormal glucose metabolism, cardiomyopathy, hypercholesterolemia, hypertriglyceridemia, hypertension, myocardial infarction, and obesity. Only incident CHCs that occurred after the blood draw for methylation profiling were considered.

### 4.2 Cell Cycle Modeling

We adopted the following mathematical model from Pugh et al [13] to model the cell cycle:

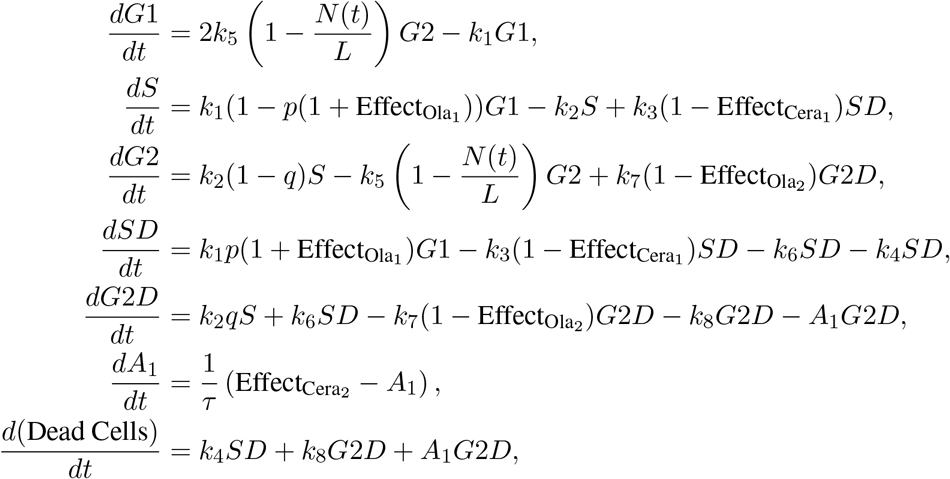

where *G*1, *S*, and *G*2, represent the density of cells over time that are in the corresponding stages of the cell cycle. Furthermore, *SD* and *G*2*D* are cells that are damaged and arrested at the S *→* G2 or G2 *→* M checkpoints, and *A*_1_ is a transit compartment to describe cell death, where Dead Cells is the total number of dead cells. The total cell density in the system is given by *N* (*t*) = *G*1 + *S* + *SD* + *G*2 + *G*2*D* + Dead Cells and *L* is the carrying capacity. Drug effects are modeled using the following sigmoid Emax equations:

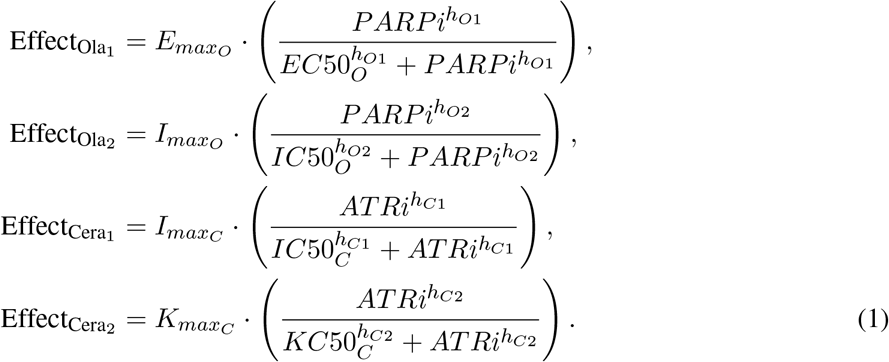

All parameter values were adopted from Pugh et al [13] and are listed in Table 2.

**Table 2:** Parameters used in cell-cycle model.

| Parameter | Value | Definition |
| --- | --- | --- |
| $k_1$ | 0.050779 | Transition rate from G1 |
| $k_2$ | 0.17906 | Transition rate from S |
| $k_3$ | 0.39181 | Rate of repair of SD cell |
| $k_4$ | 1.1443 | Death rate of SD cell |
| $k_5$ | 0.060188 | Transition rate from G2/M |
| $k_6$ | 0.32358 | Transition rate from SD to G2D after incorrect repair |
| $k_7$ | 2.1526 | Rate of repair of G2D cell |
| $k_8$ | 0.0010523 | Death rate of G2D cell |
| $p$ | 0.14826 | Probability that cell will enter SD from G1 |
| $q$ | 0.45458 | Probability that cell will enter G2D from S |
| $L$ | 13727.7261 | Carrying capacity |
| $\tau$ | 95.9914 | Time spent in death delay |
| $I_{max_C}$ | 0.92714 | Maximal inhibitory effect of ATRi |
| $IC50_C$ | 5.3318 | ATRi concentration resulting in half the maximal inhibitory effect |
| $h_{C1}$ | 5.1795 | Hill coefficient |
| $K_{max_C}$ | 2.8866 | Maximal killing effect of ATRi |
| $KC50_C$ | 0.14143 | ATRi concentration resulting in half the maximal killing effect |
| $h_{C2}$ | 5.6103 | Hill coefficient |
| $E_{max_O}$ | 1.9721 | Maximal effect of PARPi |
| $EC50_O$ | 0.15942 | PARPi concentration resulting in half the maximal effect |
| $h_{O1}$ | 0.40941 | Hill coefficient |
| $I_{max_O}$ | 1 | Maximal inhibitory effect of PARPi |
| $IC50_O$ | 0.041212 | PARPi concentration resulting in half the maximal inhibitory effect |
| $h_{O2}$ | 0.10945 | Hill coefficient |
| $G1_0$ | 687 | Initial number of G1 cells |
| $S_0$ | 199.5 | Initial number of S cells |
| $G2_0$ | 586.5 | Initial number of G2 cells |
| $SD_0$ | 23.55 | Initial number of SD cells |
| $G2D_0$ | 3.45 | Initial number of G2D cells |
| Dead Cells<sub>0</sub> | 0 | Initial number of dead cells |
| $A1_0$ | 0 | Initial number of cells in transit compartment |

### 4.3 Microbiome Modeling

We have developed a system of ordinary differential equations (ODE) to explore how chemotherapy’s impact on the microbiome affects the endogenous immune system and response to CER T cell therapy. We consider seven main state variables in our model: i) tumor cells (*C*(*t*)), ii) immune cells (*I*(*t*)), iii) the plasticity function (*P* (*t*)) capturing whether the immune environment is anti-tumor (*P* = 0) or pro-tumor (*P* = 1), iv) the levels of SCFAs (*S*(*t*)) produced by the (healthy) gut microbiome which mediate the immune state, *P*, v) the carrying capacity of SCFAs (*K*(*t*)) representing their steady state levels, vi) the concentration of chemotherapy (*D*(*t*)), and vii) the density of CER T-cells (*T* (*t*)). This leads to a system of coupled ODEs, where *α* values indicate growth rates, *κ* values are consumption rates, *γ* is used for death/kill rates from interactions, and *δ*values indicate intrinsic death/decay rates (see Table 3 for details):

**Table 3:** Parameters used in microbiome model.

| Parameter | Value | Definition |
| --- | --- | --- |
| $\alpha_S$ | 0.4 | Base growth rate of SCFAs |
| $\alpha_T$ | 0.1 | Base growth rate of CER T cells |
| $\alpha_F$ | 0.03 | Growth rate increase due to SCFAs |
| $\alpha_P$ | 0.02 | Growth rate change due to immune cell plasticity |
| $\alpha_I$ | 0.45 | Base growth rate of immune cells |
| $\alpha_C$ | 0.05 | Base growth rate of cancer cells |
| $\kappa_I$ | 0.01 | Consumption rate of SCFAs by immune cell |
| $\kappa_T$ | 0.01 | Consumption rate of SCFAs by CER T cells |
| $\gamma_{DS}$ | 0.5 | Destruction rate of SCFAs due to the drug |
| $\gamma_{DT}$ | 0.95 | Death rate of immune & CER T cells due to the drug |
| $\gamma_D$ | 0.5 | Death rate of cancer cells due to the drug |
| $\gamma_I$ | 0.03 | Death rate of cancer cells due to immune cells |
| $\gamma_T$ | 0.08 | Death rate of cancer cells due to CER T cells |
| $\delta_D$ | 0.2 | Decay rate of the drug |
| $\delta_T$ | 0.37 | Decay rate of CER T cells |
| $\delta_K$ | 0.001 | Decay rate of the long term SCFA reduction |
| $D_d$ | 1.0 | Drug dose |
| $K_d$ | $0.1D_dS_0$ | Decrease in SCFA carrying capacity due to drug application |
| $S_0$ | – | Initial concentration of SCFAs |

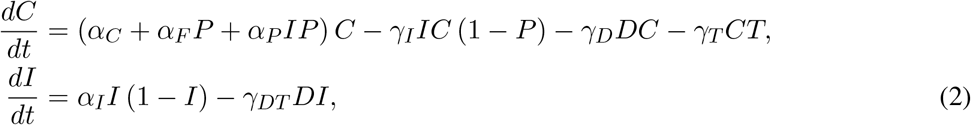

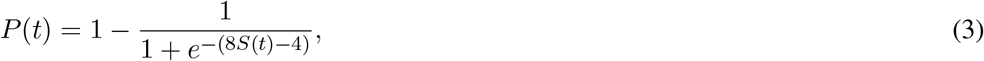

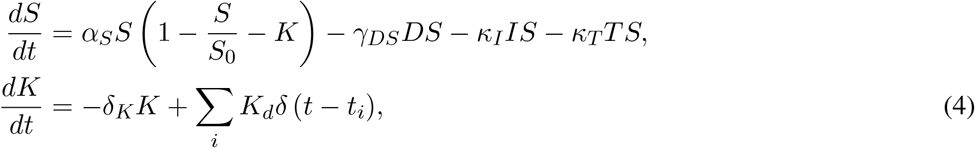

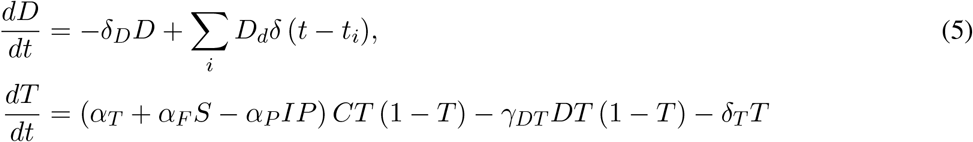

In short, we assume that tumor cells (*C*(*t*)) grow exponentially and are killed by immune cells (*I*(*t*)), drug (*D*(*t*)), and CER T-cells (*T* (*t*)) in a way that is modulated by the immune state *P* (*t*), which in turn is driven by the ratio of long to short chain fatty acids. Specifically, we assume that SCFA levels, *S*, are proportional to microbiome population size, and increase logistically until carrying capacity, *K*. We model the effect of chemotherapy as reducing not only tumor size, but also the number of immune cells (*I*), CER T-cells (*T*), and the microbiome health, thereby promoting an immunosuppressive immune response.

Specifically, we assume that each cycle of chemotherapy reduces the SCFA producers’ steady state level (*K*(*t*)), and while this level recovers over time, it does so slowly. As *P* shifts towards an immunosuppressive state (*P* = 1) due to low SCFA levels (*S*), the effect is a reduction of CER T cell (*T*) growth and an increase in proliferation rate of the tumor (*C*). As the immunosuppressive environment inhibits CER T, the result is reduced CER T attack on cancer cells, *C*. However, as SCFA increase, *P* shifts towards a pro-inflammatory/anti-tumor state (*P* = 0), promoting CER T function and increase tumor kill.

## Acknowledgments

We would like to thank the Moffitt Cancer Center Department of Integrated Mathematical Oncology (IMO) Chair, Dr. Alexander Anderson, for organizing the 12th Annual Moffitt IMO workshop: Toxicity, where this project was conceived. We are also extremely grateful to the Moffitt Cancer Center and the Moffitt PSOC for supporting this workshop through the NCI U54CA193489 grant. Finally, we would like to thank Mrs Danae Paris for her tireless work behind the scenes that allows the workshop to happen.

